# A Longitudinal Study of Antimicrobial Resistance in *Enterococcus* spp. Isolated from a Beef Processing Plant and Retail Ground Beef

**DOI:** 10.1101/2021.05.28.446255

**Authors:** Devin B. Holman, Cassidy L. Klima, Katherine E. Gzyl, Rahat Zaheer, Cara Service, Tineke H. Jones, Tim A. McAllister

## Abstract

Antimicrobial use in food-producing animals has come under increasing scrutiny due to its potential association with antimicrobial resistance (AMR). Monitoring of AMR in indicator microorganisms such as *Enterococcus* spp. in meat production facilities and retail meat products can provide important information on the dynamics and prevalence of AMR in these environments. In this study, swabs or samples were obtained from various locations in a commercial beef packing operation (n = 600 total) and from retail ground beef (n = 60) over a 19-month period. All samples/swabs were enriched for *Enterococcus* spp. and suspected enterococci isolates were identified using species-specific PCR primers. *Enterococcus faecalis* was the most frequently isolated species followed by *Enterococcus hirae,* which was found mostly on hides and ground beef. *Enterococcus faecium* (n = 9) and *E*. *faecalis* (n = 120) isolates were further characterized for antimicrobial resistance and resistant genes due to the clinical significance of these species. Twenty-one unique AMR profiles were identified, with 90% of isolates resistant to at least two antimicrobials, and two that were resistant to nine antimicrobials. Tetracycline resistance was observed most often in *E*. *faecalis* (28.8%) and was likely mediated by *tet*(M). Genomic analysis of selected *E*. *faecalis* and *E*. *faecium* isolates revealed that many of the isolates in this study clustered with other publicly available genomes from ground beef, suggesting that these strains are well adapted to the beef packaging environment.

**IMPORTANCE:** Antimicrobial resistance (AMR) is a serious challenge facing the agricultural industry. Understanding the flow of antimicrobial resistant-bacteria through the beef fabrication process and into ground beef is an important step in identifying intervention points for reducing AMR. In this study we used enterococci as indicator bacteria for monitoring AMR in a commercial beef packaging facility and in retail ground beef over a 19-month period. Although washing of carcasses post-hide removal reduced the isolation frequency of *Enterococcus* spp., a number of antimicrobial resistant-*Enterococcus faecalis* isolates were recovered from ground beef produced in the packaging plant. Genome analysis showed that several *E*. *faecalis* isolates were genetically similar to publicly available isolates recovered from retail ground beef in the United States.

## INTRODUCTION

*Enterococcus* spp. are often used as indicators of fecal contamination due to their association with the mammalian gastrointestinal tract and persistence in the environment (1). The concentration of enterococci in the feces of cattle varies, but is typically around 10^4^ to 10^5^ CFU g^−1^ (2, 3). Previous studies have reported that *Enterococcus* spp. are prevalent in ground beef samples in North America (4–7) but less information is available regarding the prevalence of enterococci in the beef processing environment.

Presently, there are more than 60 species of *Enterococcus* and two subspecies (LPSN; http://www.bacterio.net) with *Enterococcus faecalis* and *Enterococcus faecium* most frequently associated with ground beef (4, 5). Certain strains of these species are also responsible for serious nosocomial infections and vancomycin-resistant enterococci (VRE) strains are particularly difficult to treat (8, 9). Many enterococci are intrinsically resistant to several antimicrobials and also acquire resistance through horizontal gene transfer and point mutations (10, 11).

Feedlots in North America have traditionally administered antimicrobials to cattle to prevent and treat disease (12). This includes classes of antimicrobials that are also used in human medicine such as β-lactams, fluoroquinolones, macrolides, and tetracyclines (13, 14). However, there is concern that the use of antimicrobials in food-producing animals selects for antimicrobial-resistant bacteria that may be disseminated to humans through food and the environment (15). Resistant strains of *E*. *faecium* isolated from meat have colonized the human GI tract in challenge experiments (16) and transfer of the tetracycline resistance gene, *tet*(M) from an *E*. *faecium* strain of meat origin to human clinical enterococci isolates has been demonstrated *in vitro* (17). The culturability and ubiquity of *Enterococcus* spp. in cattle make them ideal for monitoring antimicrobial resistance (AMR) in beef processing facilities and retail products.

Therefore, in this study we isolated enterococci from samples taken from a commercial beef processing facility over a nineteen-month period and from retail ground beef. The antimicrobial susceptibility of selected *E*. *faecalis* and *E*. *faecium* isolates was determined and a subset of these isolates were further characterized using whole genome sequencing. These sequenced genomes were also compared with publicly available *E*. *faecalis* and *E*. *faecium* genomes from different sources.

## RESULTS

### Enterococcus spp. distribution and prevalence

Ten different *Enterococcus* species were isolated from swabs and ground beef samples with *E*. *faecalis*, *Enterococcus hirae*, and *E*. *faecium* most frequently recovered (Table 1). Within the beef processing facility, the carcasses after hide removal and the ground beef yielded the greatest number of samples positive for enterococci. *E*. *faecalis* was the only species from all five sampling locations.

**Table 1.** Distribution and prevalence of *Enterococcus* spp. in swabs and samples from four different locations in a beef processing facility (n = 150) and in retail ground beef (n = 60). Values represent the number of positive swabs or samples and include isolates from both selective (erythromycin) and non-selective media.

|  | Hide<br>removal | After<br>final<br>washing | Conveyor<br>belt | Ground<br>beef from<br>processing<br>facility | Ground<br>beef from<br>retail | Total |
| --- | --- | --- | --- | --- | --- | --- |
| *E. faecalis* | 32 | 11 | 11 | 114 | 41 | **209** |
| *E. hirae* | 45 | 3 | 0 | 30 | 0 | **78** |
| *E. faecium* | 4 | 2 | 0 | 5 | 7 | **18** |
| *E. raffinosus* | 0 | 0 | 1 | 8 | 0 | **9** |
| *E. malodoratus* | 2 | 2 | 2 | 2 | 0 | **8** |
| *E. durans* | 8 | 0 | 0 | 0 | 0 | **8** |
| *E. gallinarum* | 1 | 0 | 2 | 0 | 1 | **4** |
| *E. casseliflavus* | 3 | 0 | 0 | 0 | 0 | **3** |
| *E. avium* | 0 | 0 | 0 | 0 | 1 | **1** |
| *E. mundtii* | 1 | 0 | 0 | 0 | 0 | **1** |
| % Positive for<br>*Enterococcus*<br>spp. | 51.3% (77) | 12.0%<br>(18) | 10.7% (16) | 82.7%<br>(124) | 81.7% (49) |  |

### Antimicrobial susceptibility and detection of antimicrobial resistance genes

Antimicrobial susceptibility testing was done on 120 *E*. *faecalis* and 9 *E*. *faecium* isolates using 16 different antimicrobials (Table S1). Nearly all *E*. *faecalis* isolated on non-selective media were resistance was to lincomycin (97.4%) and quinupristin-dalfopristin (92.8%) (Table 2). Phenotypic resistance to ciprofloxacin (10.8%), erythromycin (9.0%), and tetracycline (28.8%) was also noted in several *E*. *faecalis* isolates. Although there were fewer *E*. *faecium* isolates available for testing, resistance phenotypes were similar to *E*. *faecalis* with the exception of ciprofloxacin resistance, which was not observed in any of the *E*. *faecium* strains. Two *E*. *faecalis* isolates (H11 and H22) from the hide removal samples were resistant to nine antimicrobials and one (G69E) from ground beef from the processing plant was resistant to six. Only one *Enterococcus* isolate was susceptible to all 16 antimicrobials tested; however, no resistance was recorded for linezolid, penicillin, or vancomycin for any of the isolates.

**Table 2.** Antimicrobial susceptibility for *E*. *faecalis* (n = 111) isolated on non-selective media by antimicrobial and isolation source. Values represent percentage of isolates that are resistant and numbers in parentheses indicate total number of isolates. None of the isolates were resistant to linezolid, nitrofurantoin, penicillin, tigecycline, or vancomycin.

| Antimicrobial class | Antimicrobial | Species | After hide removal (H) | After final washing (W) | Conveyor belt (C) | Ground beef from processing facility (G) | Ground beef from retail (R) | Total |
| --- | --- | --- | --- | --- | --- | --- | --- | --- |
| Aminoglycosides | GEN | *E. faecalis* | 11.1% (2) | 0 | 0 | 0 | 0 | 1.8% (2) |
|  | KAN | *E. faecalis* | 11.1% (2) | 0 | 0 | 0 | 0 | 1.8% (2) |
|  | STR | *E. faecalis* | 11.1% (2) | 0 | 0 | 0 | 0 | 1.8% (2) |
| Fluoroquinolones | CIP | *E. faecalis* | 5.6% (1) | 0 | 28.6% (2) | 11.8% (4) | 11.6% (5) | 10.8% (12) |
| Lincosamides | LIN | *E. faecalis* | 100% (18) | 100% (9) | 100% (7) | 94.1% (32) | 97.7% (42) | 97.3% (108) |
| Lipopeptides | DAP | *E. faecalis* | 0 | 0 | 0 | 5.9% (2) | 0 | 1.8% (2) |
| Macrolides | ERY | *E. faecalis* | 11.1% (2) | 11.1% (1) | 0 | 14.7% (5) | 4.6% (2) | 9.0% (10) |
|  | TYL | *E. faecalis* | 11.1% (2) | 0 | 0 | 2.9% (1) | 2.3% (1) | 3.6% (4) |
| Phenicol | CHL | *E. faecalis* | 11.1% (2) | 0 | 0 | 0 | 0 | 1.8% (2) |
| Streptogramins | SYN | *E. faecalis* | 94.4% (17) | 77.7% (7) | 100% (7) | 94.1% (32) | 93.0% (40) | 92.8% (103) |
| Tetracyclines | TET | *E. faecalis* | 11.1% (2) | 11.1% (1) | 14.3% (1) | 50.0% (17) | 25.6% (11) | 28.8% (32) |

NI: not isolated; CHL: chloramphenicol; CIP: ciprofloxacin; DAP: daptomycin; ERY: erythromycin; GEN: gentamicin; KAN: kanamycin; LIN: lincomycin; STR: streptomycin; SYN: quinupristin-dalfopristin; TET: tetracycline; TYL: tylosin.

Among the 119 *E*. *faecalis* and 9 *E*. *faecium* isolates from selective and non-selective media displaying phenotypic resistance to at least one antimicrobial, there were 21 unique AMR profiles (Table S2). The most common AMR profiles included resistance to quinupristin-dalfopristin and lincomycin (52.3%; 67) and quinupristin-dalfopristin, lincomycin, and tetracycline (20.3%; 26). The *E*. *faecalis* and *E*. *faecium* isolates were also screened for the presence of *erm*(B), *msrC*, *tet*(B), *tet*(C), *tet*(L), *tet*(M), *vanA*, *vanB*, and *vanC1* via PCR. The *tet*(M) (26.5%) and *erm*(B) (7.7%) genes were detected most frequently in *E*. *faecalis* and *msrC* (75.0%) and *erm*(B) (16.7%) in *E*. *faecium*. None of the *van* genes or *tet*(C) were found among these isolates.

### Genome analysis

The assembly statistics for the 47 *E*. *faecalis* and 8 *E*. *faecium* genomes sequenced are reported in Holman et al. (18) and Table S3. The size of the *E*. *faecalis* and *E*. *faecium* genomes ranged from 2,647,103 to 3,246,301 bp and 2,507,908 to 2,761,265 bp, respectively.

### Antimicrobial resistance genes within genome assemblies

We screened the *E*. *faecalis* and *E*. *faecium* assemblies for antimicrobial resistance genes (ARGs) using the CARD RGI (Comprehensive Antibiotic Resistance Database Resistance Gene Identifier) and identified 15 different ARGs conferring resistance to 8 different antimicrobial classes. Similar to the PCR-based screening of select ARGs, *tet*(M) (31.9%) and *erm*(B) (8.5%) were found most often within the *E*. *faecalis* genomes (Table 3). The genes *efrA*, *efrB*, *emeA,* and *lsa*(A) which encode for multidrug efflux pumps (19, 20) were identified in all *E*. *faecalis* genomes as was *dfrE*, a dihydrofolate reductase gene conferring resistance to diaminopyrimidine. All sequenced *E*. *faecium* genomes carried the *aac(6′)-Ii* and *msrC* genes conferring resistance to aminoglycosides and macrolides-streptogramin B, respectively. The *efmA* gene which encodes a multidrug efflux pump (21) was found in all but one of the *E*. *faecium* genomes. The *aac(6′)-Ii*, *efmA*, and *msrC* genes are considered to be intrinsic within *E*. *faecium* (10). One *E*. *faecalis* strain (H11) that had been isolated from a hide prior to washing carried 10 additional ARGs: *aac(6′)-Ie-aph(2’’)-Ia*, *aad(6)*, *ant(6)-Ia*, *aph(3′)-IIIa*, *catA8*, *erm*(B), *lsaE*, *sat4*, and *tet*(M). A different *E*. *faecalis* strain (H22) from the hides had seven additional ARGs: *aad(6)*, *ant(6)-Ia*, *aph(3′)-IIIa*, *lsaE*, *sat4*, and *tet*(M). These two isolates were phenotypically resistant to nine different antimicrobials and had the same multi-locus sequence typing (MLST) profile but were collected three months apart. The only other isolate with more than two additional ARGs, *E*. *faecalis* H96E, was also collected from hides.

**Table 3.** Antimicrobial resistance genes identified in sequenced *Enterococcus faecalis* (n = 47) and *Enterococcus faecium* (n = 8) genomes.

| Gene | Product | Target | *E. faecalis* | *E. faecium* |
| --- | --- | --- | --- | --- |
| *aac(6')-Ii* | Acetyltransferase | Aminoglycosides | 0 | 100% (8) |
| *ant(6)-Ia* | Nucleotidyltransferase | Aminoglycosides | 4.3% (2) | 0 |
| *ant(9)-Ia* | Nucleotidyltransferase | Aminoglycosides | 0 | 12.5% (1) |
| *aph(3')-IIIa* | Phosphotransferase | Aminoglycosides | 4.3% (2) | 0 |
| *lnuG* | Nucleotidyltransferase | Lincosamides | 2.1% (1) | 0 |
| *msrC* | ABC transporter | Macrolides | 0 | 100% (8) |
| *erm(A)* | 23S rRNA<br>methyltransferase | Macrolides | 0 | 12.5% (1) |
| *erm(B)* | 23S rRNA<br>methyltransferase | Macrolides | 8.5% (4) | 12.5% (1) |
| *optrA* | ABC transporter | Oxazolidinones | 0 | 12.5% (1) |
| *lpsB* | Intrinsic peptide<br>antibiotic-resistant LPS | Peptides | 2.1% (1) | 0 |
| *catA8* | Chloramphenicol<br>acetyltransferase | Phenicol | 2.1% (1) | 0 |
| *lsa(E)* | ABC transporter | Multiple drugs | 4.3% (2) | 0 |
| *sat4* | Acetyltransferase | Streptothricins | 4.3% (2) | 0 |
| *tet(45)* | Efflux protein | Tetracyclines | 2.1% (1) | 12.5% (1) |
| *tet(M)* | Ribosomal protection<br>protein | Tetracyclines | 31.9%<br>(15) | 37.5% (3) |

Three *E*. *faecalis* (H11, H22, and H96E) and two *E*. *faecium* (H112E and H134E) isolates with multidrug resistance profiles of interest were examined further to determine the genetic context of the ARGs detected. All five multidrug-resistant strains contained an insertion sequence harboring *tet*(M) (Fig. 1A) that had high sequence similarity (>80% identity and >70% coverage when aligned using *E*. *faecium* H134E) to integrative and conjugative elements found in *Streptococcus suis* (ICESsu05SC260; GenBank KX077888.1, ICESsuJH1308-2; GenBank KX077884.1). Alignment of this region in all five isolates showed 85% pairwise identity and revealed two variants with similarity in gene arrangements within *E*. *faecalis* H11, *E*. *faecalis* H22, and *E*. *faecium* H112E and between *E*. *faecium* H134E and *E*. *faecalis* H96E. Differences between the variants occurred on the left flank and included genes associated with integration and the presence of *tet*(L) [designated *tet*(45) by the CARD RGI] adjacent to *tet*(M) in H96E and H143E but not in H11, H22, and H112E. Despite complementarity, there were a significant number of point mutations in this region between H11, H22, and H112E (88% pairwise identify) that could reflect differences in the residence time of this gene region within each strain.

**Figure 1.**
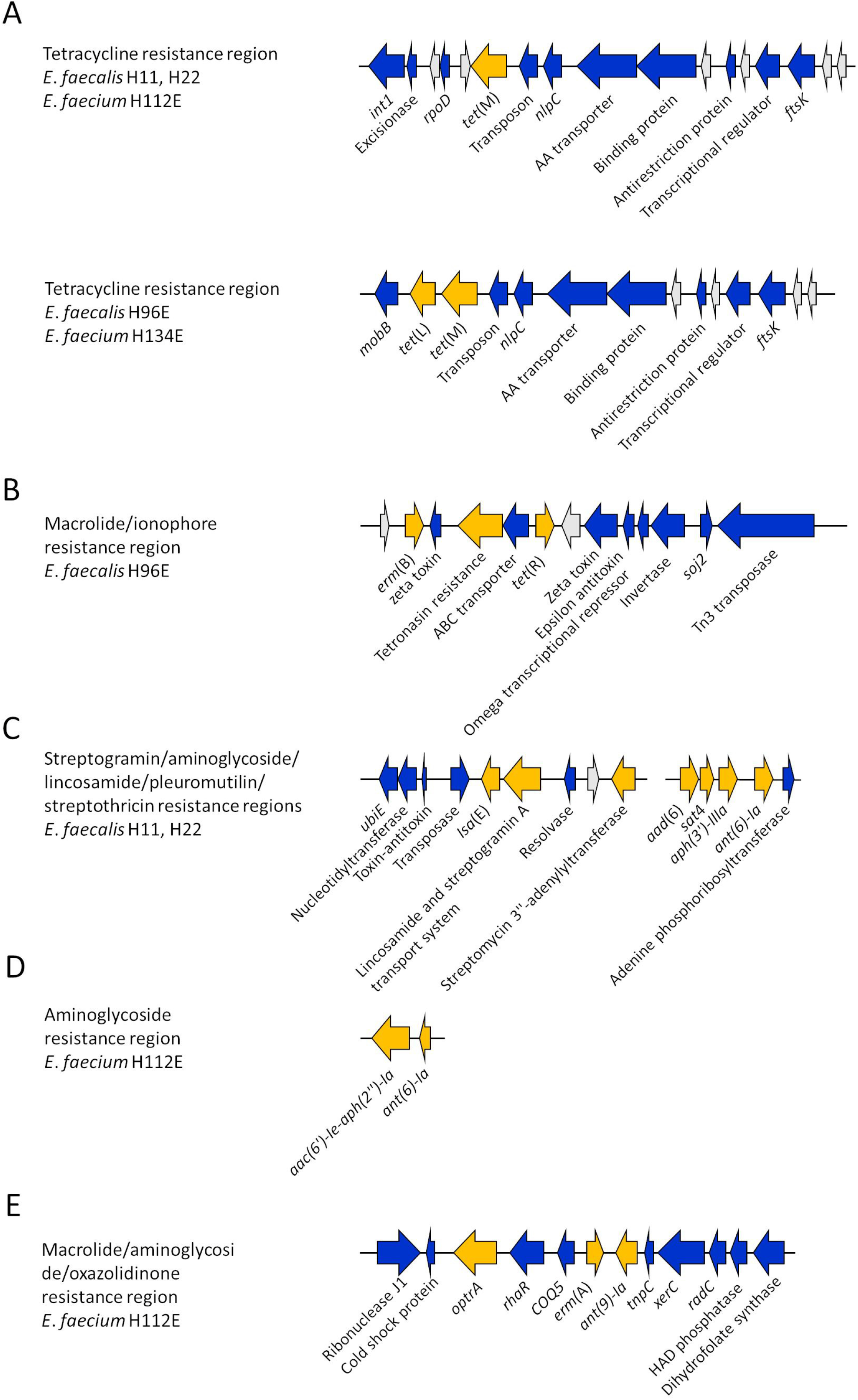
Location of antimicrobial resistance genes (ARGs) within indicated *Enterococcus faecalis* and *Enterococcus faecium* strains. The ARGs are displayed in yellow, non-ARGs genes are blue, and hypothetical proteins are colored grey.

In *E*. *faecalis* H96E, approximately 60 kb upstream of *tet*(M), *erm*(B) was found adjacent to a tetronasin resistance gene, a *tet*(R) gene, a transposase, a toxin-antitoxin system, and other genes associated with transcriptional regulation (Fig. 1B). The *erm*(B) gene was also present in *E*. *faecalis* H11 but was assembled as a single gene contig and therefore did not provide information about its location within the genome. The *lsa*(E) gene in *E*. *faecalis* H11 and H22 was found on contigs with identical gene arrangements that were truncated at the same location on the left and right flank (Fig. 1C). In addition to *lsa*(E), these contigs also contained an unnamed streptomycin 3“-adenylyltransferase and a lincosamide and streptogramin A transport system ATP-binding/permease gene. The *E*. *faecalis* H11 and H22 assemblies also had contigs carrying the *aad*(6), *sat4*, *aph(3′)-IIIa*, and *ant(6)-Ia* genes. Based on alignment against multiple *Enterococcus* strains in NCBI, the *sat4* gene-containing contig was adjacent on the chromosome to the contig carrying *lsa*(E), with the streptomycin 3′’-adenylyltransferase and *aad*(6) genes next to each other. As with other ARG regions found in these isolates, strong pairwise identity was observed between parts of these contigs and similar cassettes found in *Staphylococcus aureus* strains (*S*. *aureus* BA01611; RefSeq NC_007795.1, *S*. *aureus* MRSA_S3; RefSeq NC_007795.1).

The aminoglycoside resistance genes *aac(6′)-Ie-aph(2′’)-Ia* and *ant(6)-Ia* were found adjacent to one another, comprising a single contig in strain H11 (Fig. 1.D). This couplet of ARGs is present in many *E*. *faecium* and *E*. *faecalis* strains in NCBI, but can also be found in *Staphylococcus* spp., *Clostridium* spp., and *Campylobacter coli* strains. *E*. *faecium* H112E contained a gene region harboring the oxazolidinone resistance gene *optrA* in close proximity to the macrolide resistance gene *erm*(A), *ant(9)-Ia (*aminoglycoside resistance), and *xerC*, a tyrosine recombinase gene (Fig. 1E). This gene region aligned with complete coverage and greater than 99% identity to both a plasmid in *E*. *faecalis* (GenBank CP042214.1) and an *optrA* gene cluster in *E*. *faecium* (GenBank MK251151.1) suggesting that this gene array could have originally been a plasmid that integrated into the chromosome of *E*. *faecium* H112E. Other ARGs present that either assembled into single gene contigs or gene regions lacking other ARGs were the lincosamide resistance gene *lunG* in *E*. *faecalis* H96E, the chloramphenicol resistance gene *catA*, and *msrC* in *E*. *faecium* H134E and H112E.

### Virulence genes

Genome assemblies were also screened for virulence genes using the VirulenceFinder *Enterococcus* database. The virulence genes *ace* (collagen adhesin), *camE*, *cCF10*, *cOB1* (sex pheromones), *ebpA*, *ebpB*, *ebpC* (pili proteins), *efaAfs* (adhesion), *elrA* (enterococcal leucine rich protein A), *srtA* (sortase), *tpx* (thiol peroxidase) were found in all *E*. *faecalis* genomes (Table S4). The gelatinase-encoding *gelE* and hyaluronidase genes *hylA* and *hylB* were also detected in 74.5%, 68.8%, and 83.0% of *E*. *faecalis* genomes, respectively. Only two *E*. *faecalis* genomes carried the cytolysin genes *cylABLM* but notably these were also the strains that had the greatest number of ARGs, H11 and H22. The *efaAfm* gene, which encodes a cell wall adhesin, was found in all eight *E*. *faecium* assemblies. The *acm* gene (collagen-binding protein) was the only other virulence gene detected in the *E*. *faecium* genomes (75%).

### Phylogeny of enterococcal strains

Phylogenetic relationships among the 47 *E*. *faecalis* and 8 *E*. *faecium* strains and several publicly available *E*. *faecalis* and *E*. *faecium* genomes were determined using the core genes within each species. These additional *E*. *faecalis* and *E*. *faecium* strains included all publicly available isolates from ground beef and several randomly selected human and cattle fecal isolates also from Alberta (22). The core genome of *E*. *faecalis* contained 1,325 genes and the pangenome 9,558. Among the 27 *E*. *faecium* genomes included for analysis, there were 1,417 genes in the core genome and 7,848 in the pan-genome.

*E*. *faecalis* strains clustered by MLST type (Fig. 2). Interestingly, certain *E*. *faecalis* strains that had been collected from retail ground beef in the United States had a MLST profile (ST192, ST228, and ST260) that was shared with strains isolated from the conveyor belt, carcasses after final washing, and retail ground beef in the present study. Six of the *E*. *faecalis* isolates (G92, G127E, G149, H4, W97, and W133) had the same MLST profile as one of the Alberta human isolates (HC_NS0077; leg wound). However, it should be noted that this human isolate carried *tet*(M) and an additional virulence gene which was absent from the six isolates.

**Figure 2.**
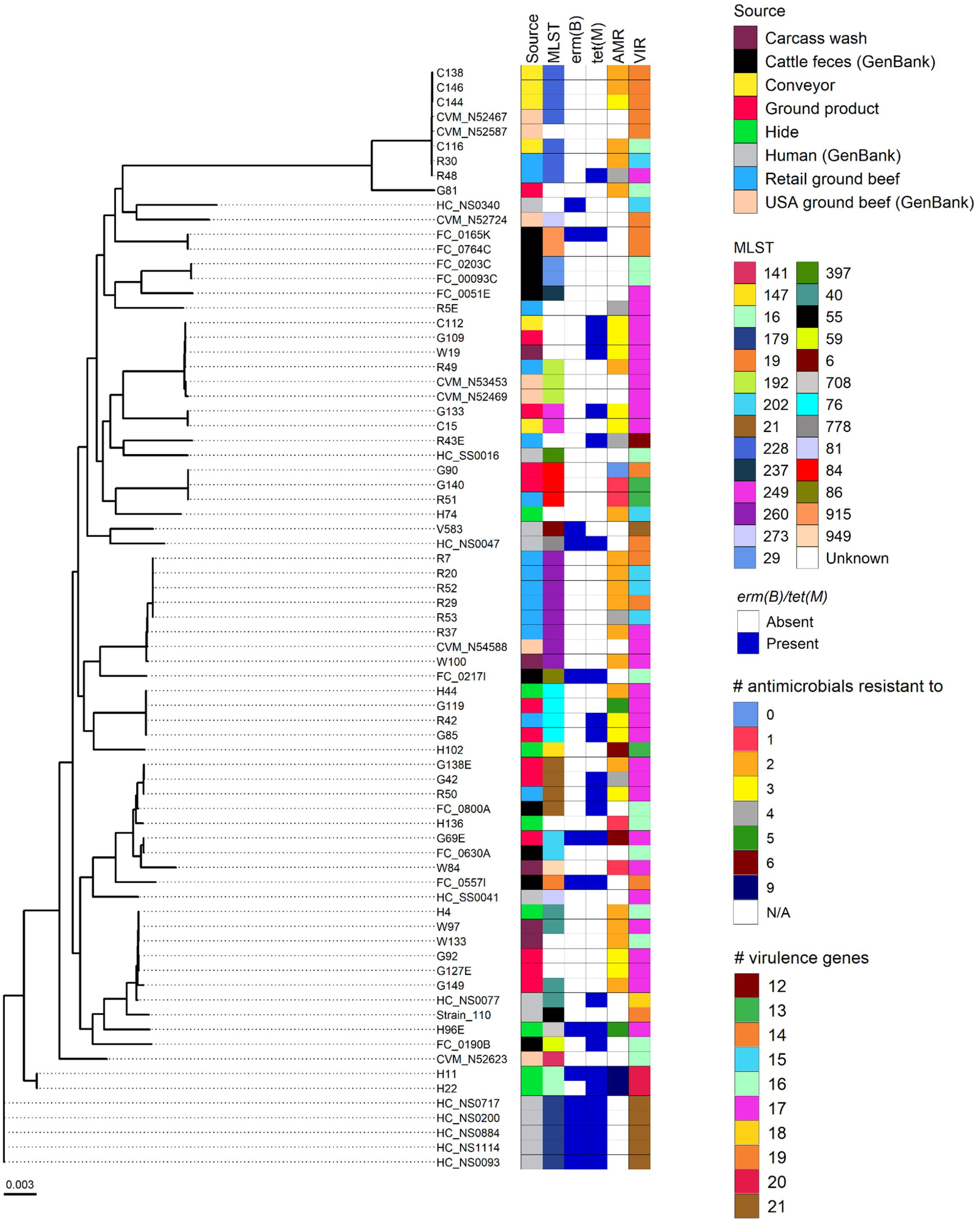
Maximum likelihood phylogeny of 47 *Enterococcus faecalis* isolates from the current study and selected publicly available *E*. *faecalis* genomes from cattle feces (n = 10), ground beef (n = 7), and humans (n = 12). Phylogeny was inferred from the alignment of 1,325 core genes using RAxML. Scale bar represents substitutions per nucleotide.

*E*. *faecium* isolates also clustered by MLST (Fig. 3). Three *E*. *faecium* isolates from retail ground beef along with two isolates from the post-wash carcasses and one from US ground beef had the same MLST (ST76). Unlike the *E*. *faecalis* genomes, there also appeared to be two distinct clades of *E*. *faecium* with the two hide isolates (H134E and H112E) in a separate clade from the other *E*. *faecium* isolates examined.

**Figure 3.**
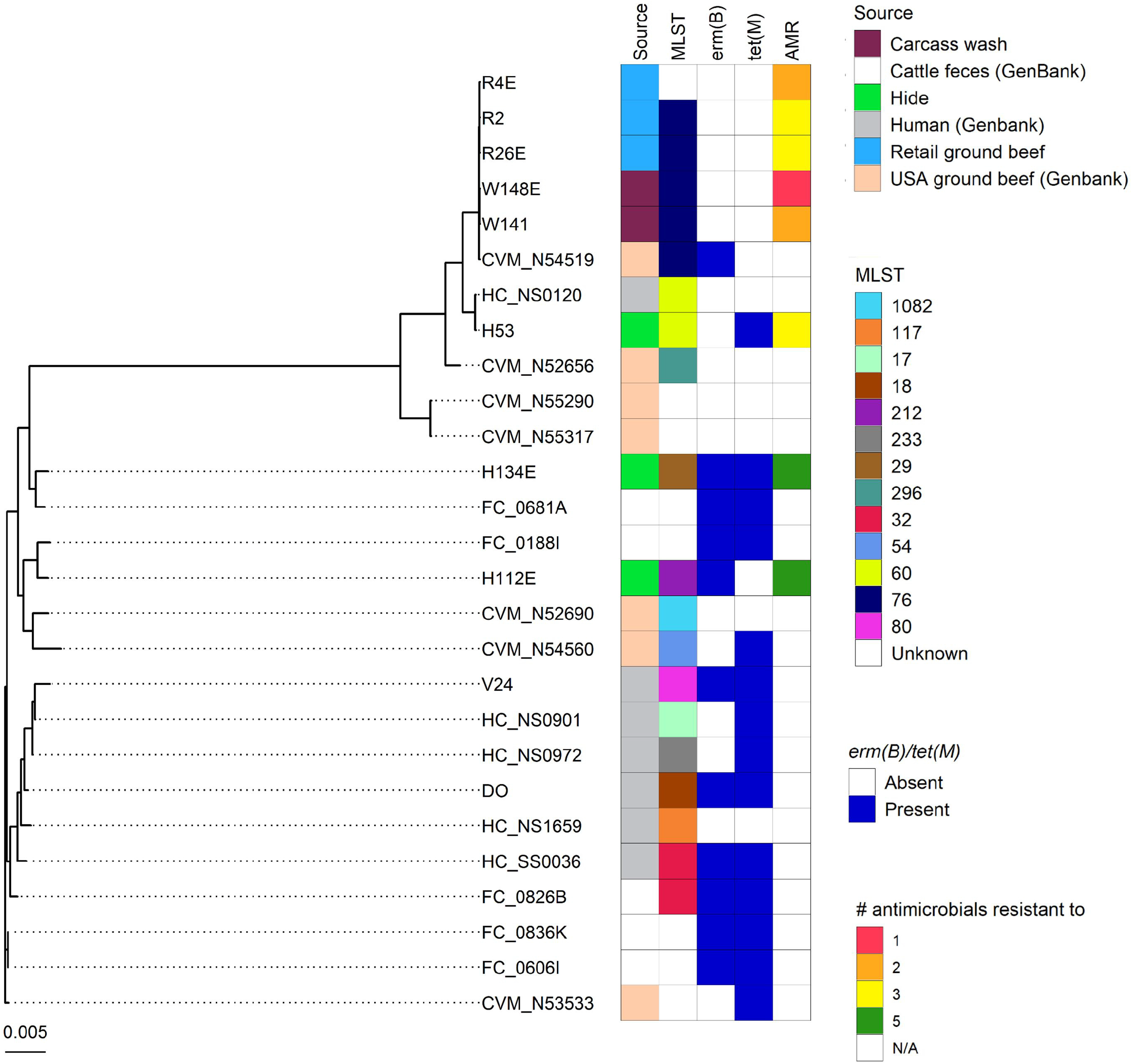
Maximum likelihood phylogeny of 8 *Enterococcus faecium* isolates and selected publicly available *E*. *faecium* genomes from cattle feces (n = 5), ground beef (n = 7), and humans (n = 7). Phylogeny was inferred from the alignment of 1,417 core genes using RAxML. Scale bar represents substitutions per nucleotide.

## Discussion

Antimicrobial resistance continues to be a serious public health threat and there are concerns that antimicrobial-resistant bacteria in food-producing animals may be transferred to humans through the food production system. In this study we used culturing and whole genome sequencing to monitor AMR and enterococci distribution in beef production from slaughter through to the retail sector over a nineteen-month period. Although 10 different *Enterococcus* spp. were isolated at least once during the study, only *E*. *faecalis* was found in all sampling locations. This is consistent with previous surveys that sampled from beef plants (4) or retail ground beef (5). *E*. *hirae* was isolated most frequently from post-hide removal swabs, which was expected given that *E*. *hirae* has been reported to be the most prevalent *Enterococcus* spp. in cattle feces (2, 22, 24) and there is greater likelihood of contamination from feces at the hide removal step (25).

The number of enterococci-positive samples recovered from the carcass post-washing and the conveyor belt area was substantially lower than in any other sample type. Carcasses are subjected to washing with hot water and spraying with organic acids after hide removal which reduces the microbial load on the carcasses. The proportion of enterococci isolated from the conveyor belts was lower than an earlier study at the same plant (10.7% vs. 48%) (4). This may represent improvements in sanitation within the conveyor area or possibly variation in the prevalence of enterococci. However, 82.7% of the ground beef produced within the plant was positive for *Enterococcus* spp., most of which were *E*. *faecalis,* suggesting that the conveyor area is not a reflection of the prevalence of enterococci in the ground beef produced. Enterococci were also isolated from the majority of ground beef samples taken from retail stores in Alberta which was similar to previous surveys of enterococci in retail ground beef in Alberta (4, 26) and the United States (5).

We subjected 120 *E*. *faecalis* and 9 *E*. *faecium* isolates to antimicrobial susceptibility testing due to their relevance to human health. Of the antimicrobials classified by the World Health Organization (WHO) as critically important in human medicine (27), infrequent resistance to ciprofloxacin, daptomycin, erythromycin, gentamicin, kanamycin, and tigecycline was noted. None of the isolates were resistant to vancomycin or linezolid, antimicrobials often used to treat VRE strains (28). Resistance to lincomycin and quinupristin-dalfopristin is intrinsic in *E*. *faecalis* and mediated by the chromosomally-encoded *lsa*(A) gene (29), thus explaining the widespread resistance of *E*. *faecalis* to these antimicrobials. Tetracycline resistance was observed in 30% of *E*. *faecalis* and 33.3% of *E*. *faecium* isolates, which may have been due to the *tet*(M) gene which was detected in 83.3% of tetracycline-resistant *E*. *faecalis* isolates and was absent in tetracycline-susceptible ones. Feedlot cattle in Western Canada have historically received tetracyclines such as chlortetracycline and oxytetracycline in feed or via injection for treatment and prevention of disease, possibly accounting for the prevalence of tetracycline resistance noted here (13).

Ionophores are one of the most widely used classes of antimicrobials in livestock production. Because they are only employed in veterinary medicine it is assumed that their use does not impact human health (30). As a potential human pathogen that inhabits the gastrointestinal tract of food-producing animals, several studies have examined ionophore resistance in *Enterococcus* spp. but reported little or no concern for its development (31). If any degree of resistance was observed it was attributed to thickening of the cell wall, or glycocalyx; traits that were considered to be genetically unstable and reversible upon removal of selective pressure (32). Recently, enterococci isolated from various locations around the world and from both humans and animals, contained both the narasin gene which encodes for ionophore-resistance and the *vanA* gene, raising the possibility that ionophore use may co-select for vancomycin resistance in these strains (30). The existence of an isolate harboring both *erm*(B) and a tetronasin resistance gene in our study merits further work to investigate possible linkages between the use of in-feed ionophores and macrolide resistance.

A large portion of the ARG cassettes examined here are also found in *Streptococcus*, *Staphylococcus* and *Campylobacter* spp. in NCBI. Future research that examines the rates of prevalence and transmissibility of these mobile regions between and amongst these species would be of considerable value in limiting the spread of AMR in bacteria of importance in human disease. Several of the *E*. *faecalis* and *E. faecium* isolates from the post-washed carcasses, conveyor belt area, and ground beef were genetically very similar to publicly available isolates from ground beef in the United States, suggesting that these particular strains are well-adapted to the beef packaging environment.

In summary, longitudinal sampling from a commercial beef packaging facility revealed the presence of *E*. *faecalis* throughout the production environment with the greatest prevalence in ground beef produced in the plant. Other *Enterococcus* spp. were isolated infrequently or as with *E*. *hirae*, confined to the carcasses post-hide removal and ground beef in the facility. Among *E*. *faecalis* isolates, the most frequently observed non-intrinsic phenotypic antimicrobial resistance was to tetracycline, which was likely mediated through the *tet*(M) gene. Several multidrug-resistant isolates were recovered including two *E*. *faecalis* from hides which were resistant to nine different antimicrobials and carried a number of ARGs on potentially mobile elements. However, the risk that such strains found on the hides may pose to the food production system is unknown as they were not isolated in the downstream processing environment.

## Materials and methods

### Sampling and isolation of Enterococcus spp

Samples were collected a total of 15 times from July 2014 through February 2016 from a commercial beef processing facility in Alberta, Canada that processed more than 3,000 carcasses per day. During each visit 10 samples were obtained from each of four different areas within the plant: carcasses after hide removal (H), carcasses after final washing (W), conveyer belts (C), and ground beef made in the plant (G). A 2 cm × 2 cm gauze swab was used to sample a randomly selected 10 cm × 10 cm area on the surface of the carcasses and conveyor belts. In total, 150 samples were obtained from each sample type or location. During the same time period, 60 samples of retail ground beef (R) were collected from various retail locations in Alberta. The exact origin of these retail ground beef samples was unknown. All samples were transported to the lab on ice and immediately processed. The swabs and 25 g of each ground product and retail ground beef sample were transferred to a stomacher bag for homogenization and pre-enrichment with 10 ml (swabs) or 225 ml (ground product/beef) of buffered peptone water. These samples were then stomached at 260 rpm for 2 min in a Stomacher 400 Circulator (Seward, Norfolk, UK) and incubated overnight at 37°C.

One milliliter of this mixture was then added to 9 ml of Enterococcosel broth (BD, Mississauga, Ontario, Canada) with or without 8 μg ml^−1^ erythromycin (Sigma Aldrich Canada, Oakville, ON, USA) and incubated overnight at 37°C for the enrichment of enterococci. Erythromycin was chosen since macrolides are important in human and veterinary medicine and enterococci are not intrinsically resistant to this antimicrobial. Enterococcosel broth tubes displaying evidence of esculin hydrolysis (black) were streaked onto Enterococcosel agar with and without 8 μg ml^−1^ erythromycin and incubated at 37°C. After 48 h the plates were examined for colonies with black zones (esculin hydrolysis) and three colonies from each plate were re-streaked onto Enterococcosel agar and incubated for 48 h at 37°C. One positive colony from each agar plate was then transferred to 1 ml of brain heart infusion (Dalynn Biologicals, Calgary, AB, Canada) containing 15% glycerol and frozen at −80°C. Confirmation and species identification of presumptive enterococci isolates was done via PCR with the Ent-ES-211-233-F and Ent-EL-74-95-R primers (33) to amplify the *groES*-*EL* spacer region as previously described (2). *Enterococcus hirae* were identified using primers mur2h-F 5′-TATGGATACACTCGAATATCTT-3′ and 5′-ATTATTCCATTCGATTAACTGC-3′ to target the muramidase (*mur-2*) gene of *E*. *hirae* as per Zaheer et al. (22). The *groES*-*EL* amplicon from non-*E*. *hirae* isolates was sequenced on an ABI Prism 3130xl Genetic Analyzer (Thermo Fisher Scientific Inc., Mississauga, ON, Canada) to differentiate *Enterococcus* spp.

### Antimicrobial resistance screening of enterococci isolates

Due to their relevance to human health, isolates with a *groES*-*EL* spacer region that was 100% identical to *E*. *faecalis* or *E*. *faecium* were screened for antimicrobial resistance genes (ARGs) and antimicrobial sensitivity. Broth microdilution with the Sensititre NARMS Gram-positive CMV3AGPF AST plate (Trek Diagnostics, Independence, OH, USA) was used to determine the susceptibility of 120 *E*. *faecalis* and 9 *E*. *faecium* isolates to sixteen different antimicrobials. For antimicrobials in the panel, the Clinical and Laboratory Standards Institute (CLSI) or European Committee on Antimicrobial Susceptibility Testing (EUCAST) minimum inhibitory concentration (MIC) breakpoints for *Enterococcus* spp. were used to interpret the results. These isolates were also screened via PCR for the presence of the ARGs *erm*(B), *msrC*, *tet*(B), *tet*(C), *tet*(L), *tet*(M), *vanA*, *vanB*, and *vanC1* as described in Beukers et al. (2) (Table S5).

### Sequencing of selected *Enterococcus faecalis* and *Enterococcus faecalis* isolates

Forty-seven *E. faecalis* and eight *E. faecium* isolates were selected for whole genome sequencing based on their AMR profiles and sample origin. Briefly, the isolates were re-cultured from the frozen glycerol on BEA and incubated for 24 h at 37°C to obtain isolated colonies with typical morphology and colour. A single colony was then streaked onto BHI agar (Dalynn Biologicals), grown overnight at 37°C, and colonies from this plate were suspended in 10 mM Tris-1mM EDTA (TE) (pH 8.0) buffer to obtain an OD_600_ of 2.0 (2 × 10^9^ cells ml^−1^). One milliliter of this suspension was pelleted via centrifugation at 14,000 × g for 2 min. Genomic DNA was extracted from the pellet using the DNeasy Blood and Tissue kit (Qiagen, Mississauga, Ontario, Canada) with the modification that cells were incubated with agitation (150 rpm) for 45 min at 37°C in 280 μl of lysis buffer (20 mM Tris-HCl [pH 8.0], 2 mM sodium EDTA, 1.2% Triton X-100 and 20 mg ml^−1^ lysozyme) (Sigma Aldrich Canada) prior to the addition of proteinase K and 5 μl of 100 mg ml^−1^ RNase A (Qiagen). The DNA concentration was determined using a Qubit fluorometer (Thermo Fisher Scientific, Mississauga, ON, Canada). The Nextera XT DNA Library Preparation kit (Illumina Inc., San Diego, CA, USA) was used to prepare sequencing libraries that were sequenced on a MiSeq instrument (Illumina Inc., San Diego, CA, USA) with the MiSeq Reagent kit v3 (Illumina Inc.; 600 cycles) or on a NovaSeq 6000 machine (Illumina Inc.) with a SP flowcell (300 cycles).

### Genomic analysis of Enterococcus faecalis and Enterococcus faecalis isolates

Trimmomatic v. 0.39 (34) was used to remove sequencing adapters, reads with a quality score of less than 15 over a 4-bp sliding window, and reads that were less than 50 bp in length. Genomes were assembled with SPAdes v. 3.15.1 (35) in “isolate mode” and the quality of the assemblies was assessed with QUAST v. 5.0.2 (36). Potential contamination within each assembly was determined using Kraken 2 v. 2.1.1 and the minikraken2 database v. 2 (37) as well as CheckM v. 1.1.3 (38). GTDB-tk v. 1.3.0 (39) was also used to confirm the taxonomic assignments of the assemblies and Prokka v. 1.14.6 (40) was used to annotate the assemblies. Determination of MLST was done on the assembled genomes using the *E. faecalis* (https://pubmlst.org/efaecalis) and *E. faecium* (https://pubmlst.org/efaecium/) MLST databases (41, 42).

The accessory, core, and pan-genome of the *E*. *faecalis* and *E*. *faecium* genomes were identified using Roary v. 3.13.0 (43) with a BLASTp identity cut-off of ≥95%. The core genome is defined as genes present in ≥ 99% of genomes. The core genes for both species were aligned in Roary using MAFFT v. 7.475 (44) and a maximum likelihood phylogenetic tree was inferred from this alignment using RAxML v. 8.2.12 (45) and viewed with ggtree v. 2.4.1 (46) in R 3.6.1.. Several publicly available *E*. *faecalis* and *E*. *faecium* assemblies from various isolation sources, including from humans and cattle in Alberta, were also included in the core and pangenome analysis as listed in Table S6. The genome assemblies were screened for virulence genes using the VirulenceFinder 2.0 database (47) and BLASTn (≥90% identity) and for ARGs using the CARD v. 3.0.9 (48) Resistance Gene Identifier (RGI). The depicted gene regions containing ARGs were constructed and validated using contig alignments in Geneious v. 11.0.9. BLAST was used to identify highly similar regions with >80% pairwise identity in bacterial strains present in NCBI.

## ACKNOWLEDGMENTS

This project was financially supported by the Beef Cattle Research Council, Agriculture and Agri-Food Canada, the Genomics Research and Development Initiative, and the Antimicrobial Resistance-One Health Consortium of the Alberta Government Major Innovation Fund. We thank Brent Avery, Ruth Barbieri, Scott Hrycauk, Nicole Lassel, TingTing Liu, and Victoria Muehlhauser for their valuable technical assistance. We are also appreciative of bioinformatics scripts provided by Arun Kommadath.

